# Heat Shock-Induced PI(4)P Increase Drives HSPA1A Translocation to the Plasma Membrane in Cancer and Stressed Cells through PI4KIII Alpha Activation

**DOI:** 10.1101/2025.02.16.638537

**Authors:** Alberto Arce, Rachel Altman, Allen Badolian, Jensen Low, Azalea Blythe Cuaresma, Uri Keshet, Oliver Fiehn, Robert V. Stahelin, Nikolas Nikolaidis

## Abstract

HSPA1A, a major heat shock protein, is known to translocate to the plasma membrane (PM) in response to cellular stress and cancer, where it plays protective roles in membrane integrity and stress resistance. Although phosphatidylinositol 4-phosphate [PI(4)P] is essential in this translocation, the signals that trigger and facilitate HSPA1A’s movement remain undefined.Given that membrane lipid composition dynamically shifts during stress, we hypothesized that heat shock-induced PI(4)P changes are crucial for HSPA1A’s PM localization. To test this hypothesis, we investigated the mechanisms driving PI(4)P changes and HSPA1A PM localization under heat shock. Lipidomic analysis, enzyme-linked immunosorbent assay (ELISA), and confocal imaging revealed a rapid PI(4)P increase at the PM post-heat shock, with levels peaking at 0 hours and declining by 8 hours. RNA sequencing and protein quantification indicated no transcriptional increase in PI4KIII alpha, the kinase responsible for PI(4)P synthesis, suggesting an alternative regulatory mechanism. Hypothesizing that heat shock enhances PI4KIII alpha activity, we performed ELISA coupled with immunoprecipitation, confirming a significant rise in PI4KIII alpha activity following heat shock. Functional analyses further demonstrated that RNAi-mediated PI4KIII alpha depletion or pharmacological PI(4)P reduction, using GSK-A1, impairs HSPA1A’s localization to the PM, confirming that HSPA1A translocation is PI(4)P-dependent. Our findings identify PI4KIII alpha activity as a key regulator of PI(4)P accumulation and subsequent HSPA1A recruitment to the PM in stressed and cancer cells. This lipid-mediated response offers new insights into stress adaptation and potentially modifiable pathways for therapeutic interventions to control HSPA1A function in cancer.

## Introduction

Heat shock protein A1A (HSPA1A), a 70-kDa molecular chaperone, plays a critical role in the cellular stress response, maintaining protein homeostasis and promoting cell survival under adverse conditions (1,2). While primarily cytosolic, HSPA1A translocates to the plasma membrane (PM) in heat-shocked and cancer cells, where its PM-localized form (mHSPA1A) contributes to increased tumor aggressiveness, therapeutic resistance, and immune modulation. This translocation is essential for membrane stabilization, signal transduction, and other stress-adaptive functions in cancer biology and disease progression (3-10).

Despite lacking canonical lipid-binding domains, HSPA1A interacts selectively with specific anionic lipids, such as phosphatidylserine (PS) and phosphatidylinositol monophosphates (PIPs), rather than neutral lipids (6,9-19). PS has been extensively studied as a critical regulator of HSPA1A’s PM localization, driven by a combination of electrostatic and hydrophobic forces. Similarly, phosphatidylinositol 4-phosphate (PI(4)P) is another key lipid supporting HSPA1A translocation (17,19) (Figure 1). PI(4)P is known to regulate protein recruitment to membrane compartments (20), yet the mechanisms by which it facilitates HSPA1A translocation remain unexplored.

**Figure 1.**
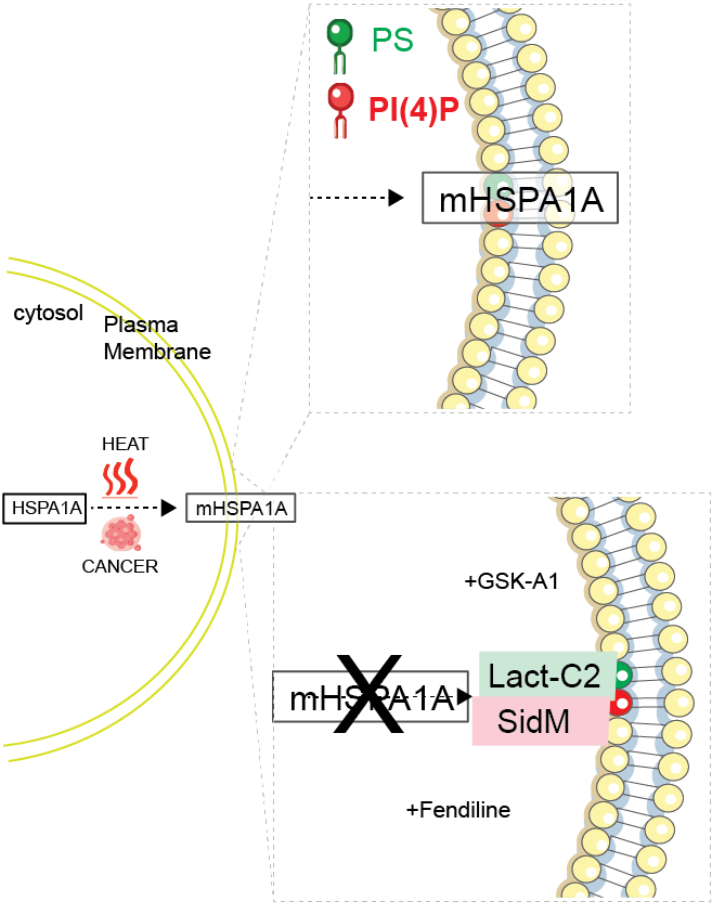
HSPA1A plasma membrane translocation is regulated by specific lipids during stress conditions. Upon heat shock or cancer-related stress, cytosolic HSPA1A translocates to the plasma membrane (PM; mHSPA1A) in a process dependent on phosphatidylserine (PS, green) and phosphatidylinositol 4-phosphate (PI(4)P, red) (14,17,19,21). The top inset highlights the PM localization of HSPA1A mediated by these lipids. Pharmacological inhibition or specific lipid-binding proteins, such as Lact-C2 (PS-binding) and SidM [PI(4)P-binding], disrupt HSPA1A translocation. GSK-A1, a PI4KIII alpha inhibitor, reduces PI(4)P levels, while fendiline interferes with PS localization, effectively preventing HSPA1A from reaching the PM.

Lipid signaling is pivotal in cellular stress responses, with changes in lipid composition and dynamics often accompanying stress and cancer states (13,15,16,22-27). Among these changes, heat stress-driven increases in specific lipids like PS and phosphatidylinositol 4-phosphate [PI(4)P] are hypothesized to play critical roles in facilitating HSPA1A’s PM localization. PS has been shown to enhance HSPA1A’s selective binding to anionic lipid-rich microdomains through electrostatic and hydrophobic forces, driving its translocation from the cytosol to the PM (3,11,14,21,28,29).

Similarly, PI(4)P—a lipid tightly regulated by PI4 kinases (PI4Ks)—is emerging as a key regulator of HSPA1A translocation (17,19). Heat shock may modulate PI(4)P levels (30-33), thereby creating PI(4)P-enriched microdomains at the PM that are crucial for HSPA1A localization. However, the mechanisms underlying this modulation remain undefined. It is unclear whether PI(4)P accumulation is driven by transcriptional or post-translational regulation of PI4Ks, changes in lipid trafficking, or other lipid-modifying processes. This gap in understanding mirrors the unanswered questions surrounding PS-mediated translocation (21), emphasizing the need to delineate the molecular triggers and pathways that regulate these lipid interactions under stress and cancer conditions.

We hypothesized that heat shock enhances PI4KIII alpha activity, leading to PI(4)P accumulation at the PM and enabling HSPA1A translocation. To test this hypothesis, we characterized cellular PI(4)P content, examined PI4KIII alpha activity, and explored the relationship between PI(4)P levels and HSPA1A localization using lipidomics, ELISA, confocal microscopy, and RNA interference (RNAi). Our findings provide new insights into lipid-mediated regulation of HSPA1A translocation and its implications for cellular stress responses and cancer progression.

## Results

### Phosphatidylinositols Increase in Response to Heat Shock

To determine whether PI(4)P levels increase after heat shock, we performed lipidomic analyses and PI(4)P quantification. Lipidomics heat maps (Figure 2A and Supplement Table 1) revealed a significant increase in the PI(4)P precursor molecule, phosphatidylinositol (PI), immediately after heat shock (0 h), followed by a reduction during recovery at 8 h, though not to baseline levels. Quantitative lipidomics data (Figure 2B) further confirmed this dynamic, indicating that PI availability increases under heat stress.

**Figure 2.**
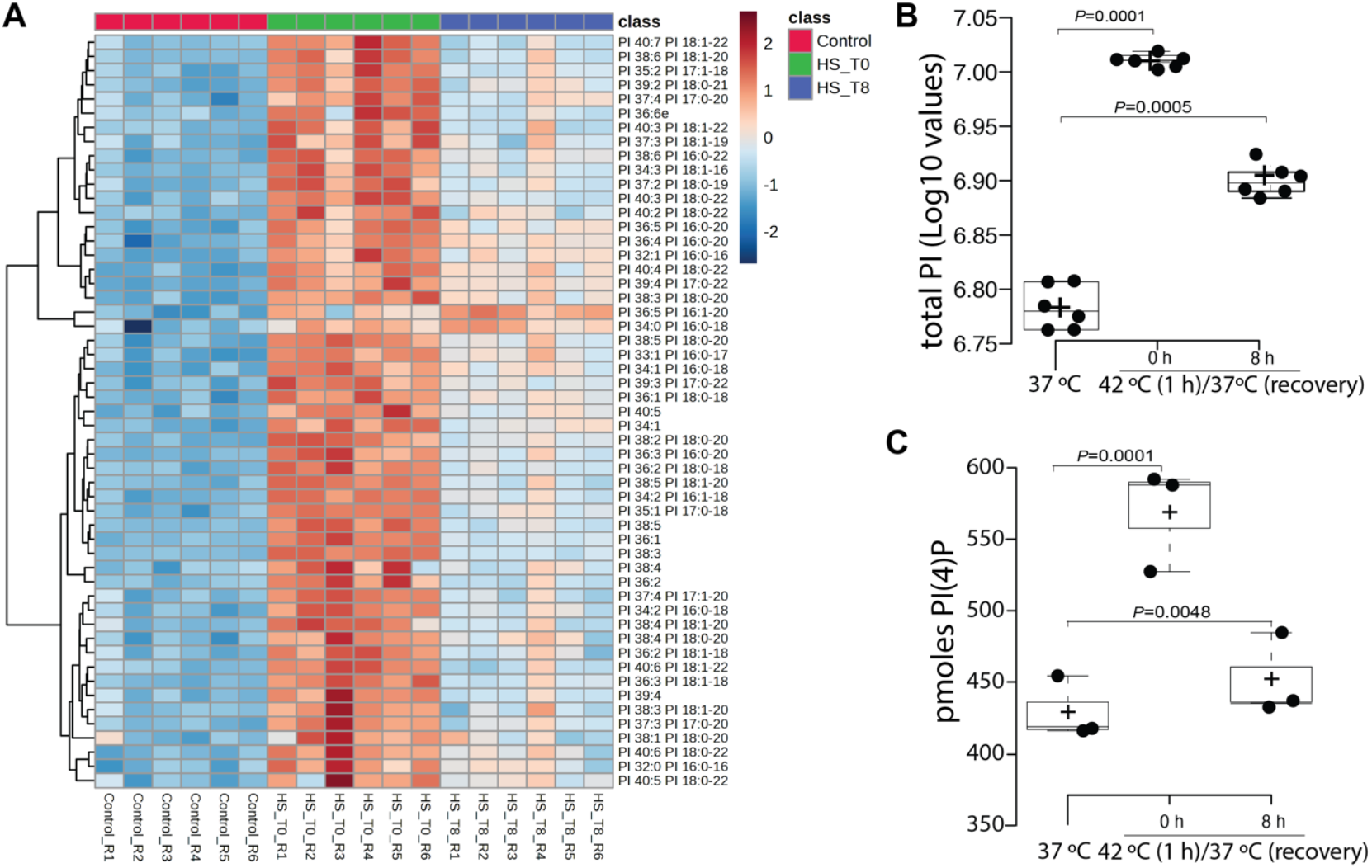
Heat shock increases phosphatidylinositol (PI) and PI(4)P levels. (A) Heatmap visualization of PI species composition in HeLa cells under control conditions (37 °C), immediately following heat shock (0 h recovery at 37 °C), and after 8 h recovery at 37 °C (8 h recovery at 37 °C). The lipid composition was calculated as the percentage of each PI subclass within the total lipidome and log-transformed for visualization. Each column represents one biological replicate. The heatmap was generated using MetaboAnalyst (34). (B) Boxplot showing total PI levels across biological replicates at 37 °C, 0 h, and 8 h post-heat shock. Total PI levels were calculated from log-transformed data of all PI species in each replicate. Center lines indicate medians; box limits represent the 25th and 75th percentiles; whiskers extend 1.5 times the interquartile range; crosses denote sample means. (C) Quantification of PI4P levels using the PI(4)P Mass ELISA Assay (Echelon Biosciences, Salt Lake, UT, USA) at multiple time points: control conditions (37 °C), immediately post-heat shock (0 h), and 8 h recovery. Data were generated using cells from three of the batches used for lipidomics.

We therefore hypothesized that PI’s increase would result in elevated PI(4)P levels. Using a PI(4)P-specific ELISA assay, we observed a sharp rise in PI(4)P levels immediately after heat shock (0 h), followed by a partial decline at 8 h recovery, yet remaining above baseline levels (Figure 2C and Supplement Figure 1). These findings suggest that heat shock induces a transient but sustained increase in PI(4)P. This dynamic increase in PI(4)P following heat shock highlights its potential role as a key signaling molecule in the cellular stress response.

### Localization and Modulation of PI(4)P at the Plasma Membrane

To determine whether PI(4)P levels increase at the PM following the observed rise in total cellular PI(4)P levels after heat shock (0 h; Figure 2C), we used confocal microscopy with the PI(4)P-specific biosensor SidM, focusing on the 0 h time point. Imaging and quantification revealed a marked increase in PI(4)P localization at the PM immediately after heat shock (0 h) (Figure 3 and Supplementary Figure 2).

**Figure 3.**
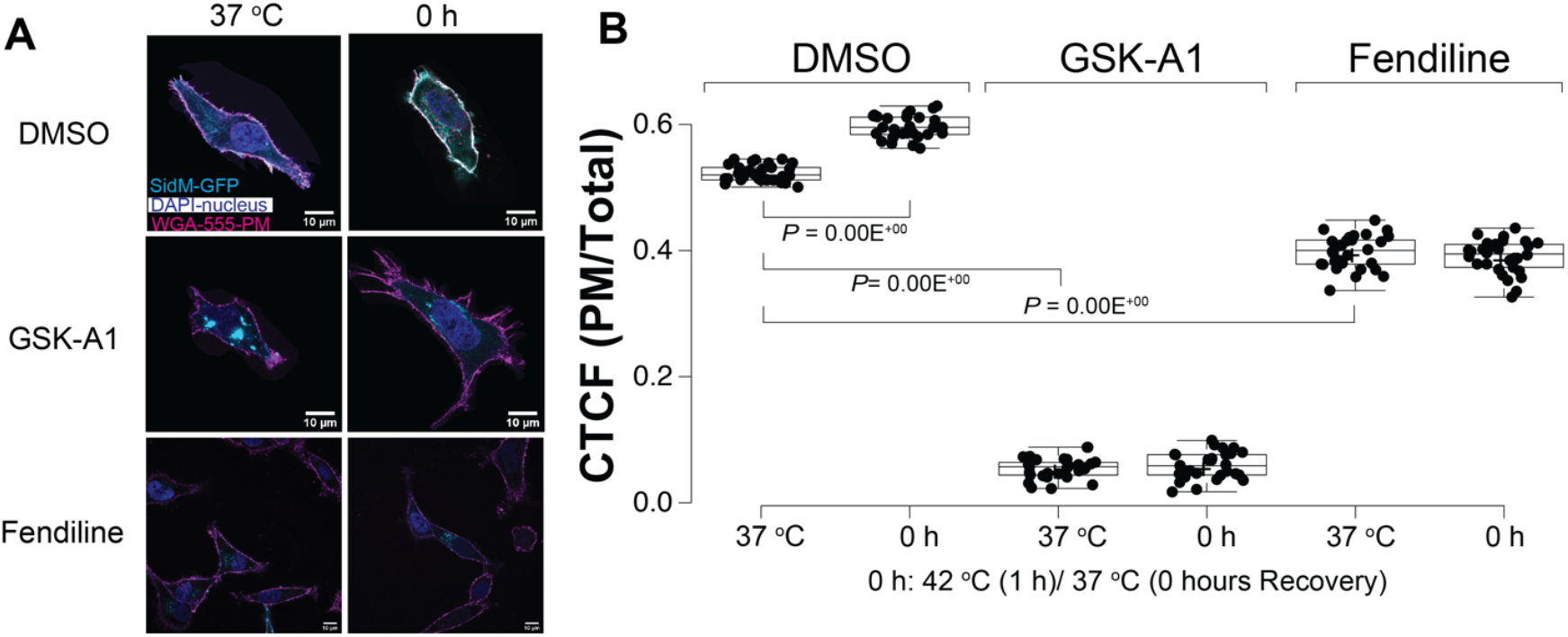
Heat shock increases PI(4)P levels at the plasma membrane (PM), and drug treatments reduce PI(4)P PM localization. (A) Representative confocal images of HeLa cells expressing SidM-P4M-GFP, a PI4P-specific biosensor, under control conditions (37 °C; left panels), immediately following heat shock (0 h recovery; right panels), and after treatment with GSK-A1 (PI4K inhibitor; middle row) or fendiline (PS modulator; bottom row). Cells were stained with WGA-FA555 (PM stain) and DAPI (nucleus stain). SidM-P4M fluorescence highlights PI4P localization at the PM. Scale bar = 10 μm. (B) Quantification of the corrected total cell fluorescence (CTCF) ratio of SidM-P4M fluorescence at the PM to the rest of the cell under control, heat-shocked, and drug-treated conditions. Data represent three independent experiments, with total individual cell counts (N = 30; shown as closed circles). Center lines indicate medians; box limits represent the 25th and 75th percentiles; whiskers extend 1.5 times the interquartile range; crosses denote sample means. Additional images are provided in Supplementary Figure 2.

To assess the impact of pharmacological inhibitors on PI(4)P distribution, we treated cells with GSK-A1 and fendiline. Both drugs reduced PI(4)P levels at the PM, with GSKA1 causing a pronounced decrease compared to the more modest effect of fendiline (Figure 3 and Supplementary Figure 2). These findings reveal the inhibitors’ specificity and the relationship between PM-localized PI(4)P and PS (35,36).

We further quantified total PI(4)P levels in the presence or absence of GSK-A1 using a PI(4)P-specific ELISA assay. Consistent with the microscopy data, total PI(4)P levels were significantly reduced in GSK-A1-treated cells, confirming its effect on PI(4)P at both the PM and cellular levels (Figure 4 and Supplement Figure 3).

**Figure 4.**
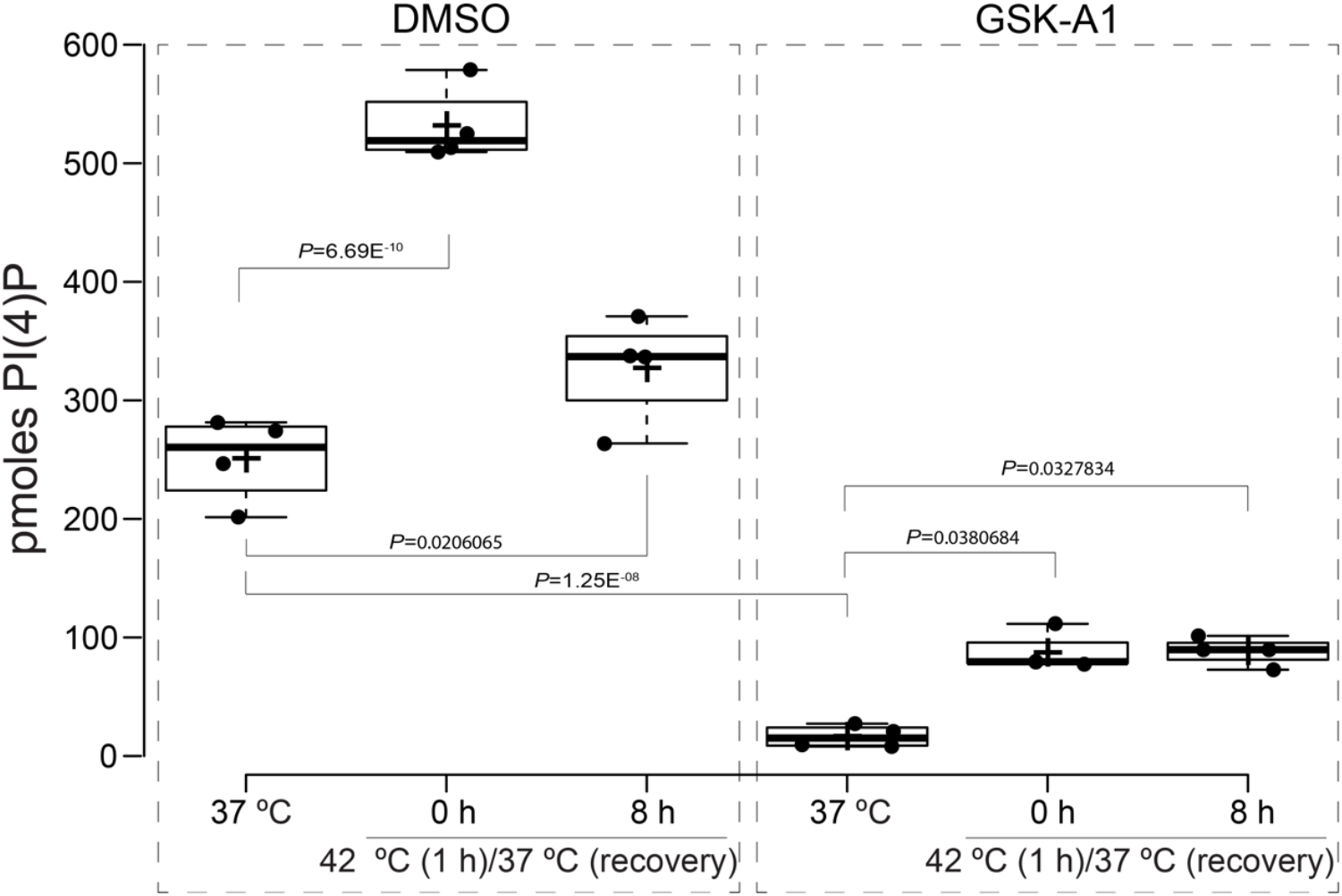
PI(4)P levels significantly increased immediately after heat shock. PI(4)P levels were measured by the PI(4)P Mass ELISA Assay (Echelon Biosciences, Salt Lake, UT, USA) under control conditions (37 °C), immediately following heat shock (0 h recovery at 37 °C), and after 8 h recovery at 37 °C. PI(4)P levels significantly increased immediately after heat shock and decreased during recovery but remained above baseline levels (left plots). Treatment with GSK-A1 (PI4K inhibitor) significantly reduced PI(4)P levels at all time points (right plots). Data represent three independent experiments, with individual data points shown as closed circles. Center lines indicate medians; box limits represent the 25th and 75th percentiles; whiskers extend 1.5 times the interquartile range; crosses denote sample means. Data were generated from different batches of cells than those used for lipidomics and other ELISA experiments in Figure 2.

These findings demonstrate that heat shock induces PI(4)P enrichment at the PM, while pharmacological inhibition with GSKA1 disrupts this localization, highlighting PI(4) P’s role in the heat shock response.

### Mechanisms Regulating PI(4)P Production

To investigate the mechanisms underlying the observed PI(4)P increase, we analyzed RNA-seq data to identify potential genes involved in PI(4)P metabolism, including those responsible for PI(4)P synthesis (e.g., PI4Ks) and modification (e.g., 5-phosphatases). A broad range of candidate genes was examined, inclduing PI4KA, PI4KB, PIP4K2A-C, and INPP5 family members. The RNA-seq analysis revealed no significant transcriptional changes correlating with PI(4)P increases. Most genes showed no change, several decreased, and only a few displayed minor increases (Figure 5 and Supplement Table 2).

**Figure 5.**
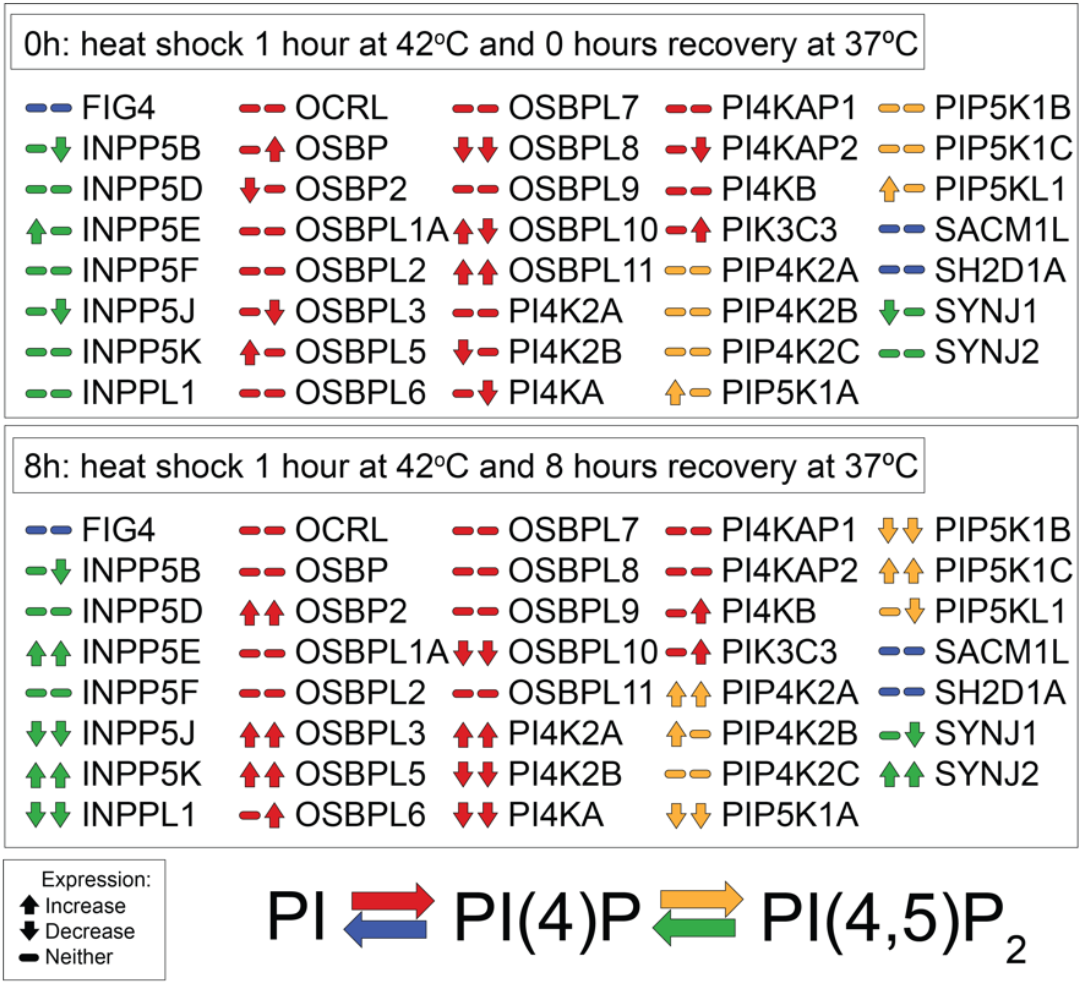
RNA-seq analysis reveals stable transcript levels and expression patterns of genes involved in PI(4)P metabolism. Schematic representation of genes associated with PI(4)P production, illustrating their expression patterns based on RNA-seq data at 0 h and 8 h post-heat shock. Each gene includes a labeled icon with two indicators reflecting results from two independent RNA-seq batches (38). Icons include upward arrows (increased expression; log2 fold change > 0.5 and adjusted p-value < 0.05), downward arrows (decreased expression; log2 fold change < -0.5 and adjusted p-value < 0.05), or dashes (no significant change). RNA-seq analysis shows that PI4KIII alpha and many other genes involved in PI(4)P metabolism exhibit stable transcript levels under heat shock conditions, indicating that transcriptional changes do not drive PI(4)P accumulation.

To verify these findings, we performed qPCR analysis for PI4KA, one of the primary PI(4)P-synthesizing enzymes, and confirmed no significant change in transcript levels (Figure 6A). We then assessed whether PI4KA protein levels increased under heat shock conditions using western blotting. The analysis showed no significant changes in total PI4KA protein levels at 0 h or 8 h post-heat shock (Figure 6B, C and Supplement Figure 4).

**Figure 6.**
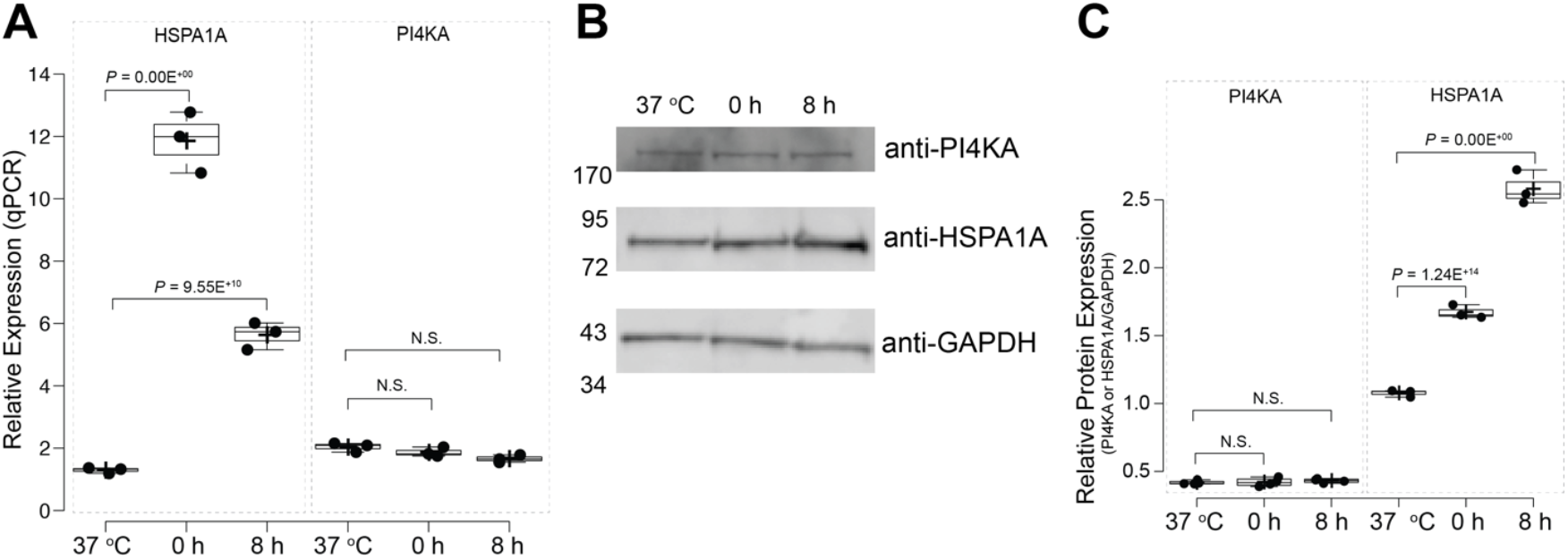
PI4KIII alpha transcript and protein levels remain stable post-heat shock. (A) qPCR analysis of PI4KIII alpha transcript stability using separate batches of cells from all other experiments, including RNA-seq. Transcript levels were assessed at control (37 °C), 0 h post-heat shock, and 8 h recovery and normalized to housekeeping genes (ACTB and GAPDH). HSPA1A was included as a positive control for heat shock response. Statistical significance was determined using one-way ANOVA (Analysis of Variance) followed by post-hoc Tukey HSD (Honestly Significant Difference) and Bonferroni tests. (B) Representative Western blot of total PI4KIII alpha protein levels in cell lysates under control conditions, with 0 h post-heat shock and 8 h recovery. HSPA1A is a positive control, demonstrating heat shock-induced upregulation, while GAPDH verifies equal protein loading (complete blots in Supplemental Figure 4). (C) Quantification of Western blot results from independent experiments showing relative PI4KIII alpha and HSPA1A protein levels normalized to GAPDH. Closed circles represent individual data points. Center lines indicate medians; box limits represent the 25th and 75th percentiles; whiskers extend 1.5 times the interquartile range; crosses denote sample means. Statistical significance was determined using one-way ANOVA (Analysis of Variance) followed by post-hoc Tukey HSD (Honestly Significant Difference) and Bonferroni tests (N.S, not significant).

These findings indicate that transcriptional or translational changes in PI4KA are not the main drivers of the observed PI(4)P increase. This observation aligns with the concept of heat shock, where non-heat shock gene expression typically decreases (37). These results suggest an alternative regulatory mechanism, likely involving an increase in enzyme activity rather than gene or protein expression changes.

### PI4KIII Alpha Activity after Heat Shock

Building on the previous results that ruled out transcriptional and translational changes in PI4KA as drivers of PI(4)P increase, we examined whether PI4KIII alpha enzyme activity was modulated by heat shock. Using the PI4-Kinase Activity ELISA kit, a competitive assay that quantifies PI(4)P produced during a kinase reaction, we assessed PI4KIII alpha activity across time points and under drug treatments.

The assay revealed a significant increase in PI4KIII alpha activity immediately after heat shock (0 h), followed by a decline during recovery at 8 h. However, activity remained above baseline levels (Figure 7A and Supplement Figures 5-6). Treatment with GSK-A1, a PI4K inhibitor, substantially reduced overall enzyme activity while preserving the heat shock-induced pattern of 0 h activation. To validate the assay, recombinant PI4KB protein was tested under identical conditions, confirming that PI(4)P production is concentration-dependent, consistent with the expected enzymatic kinetics (Figure 7B).

**Figure 7.**
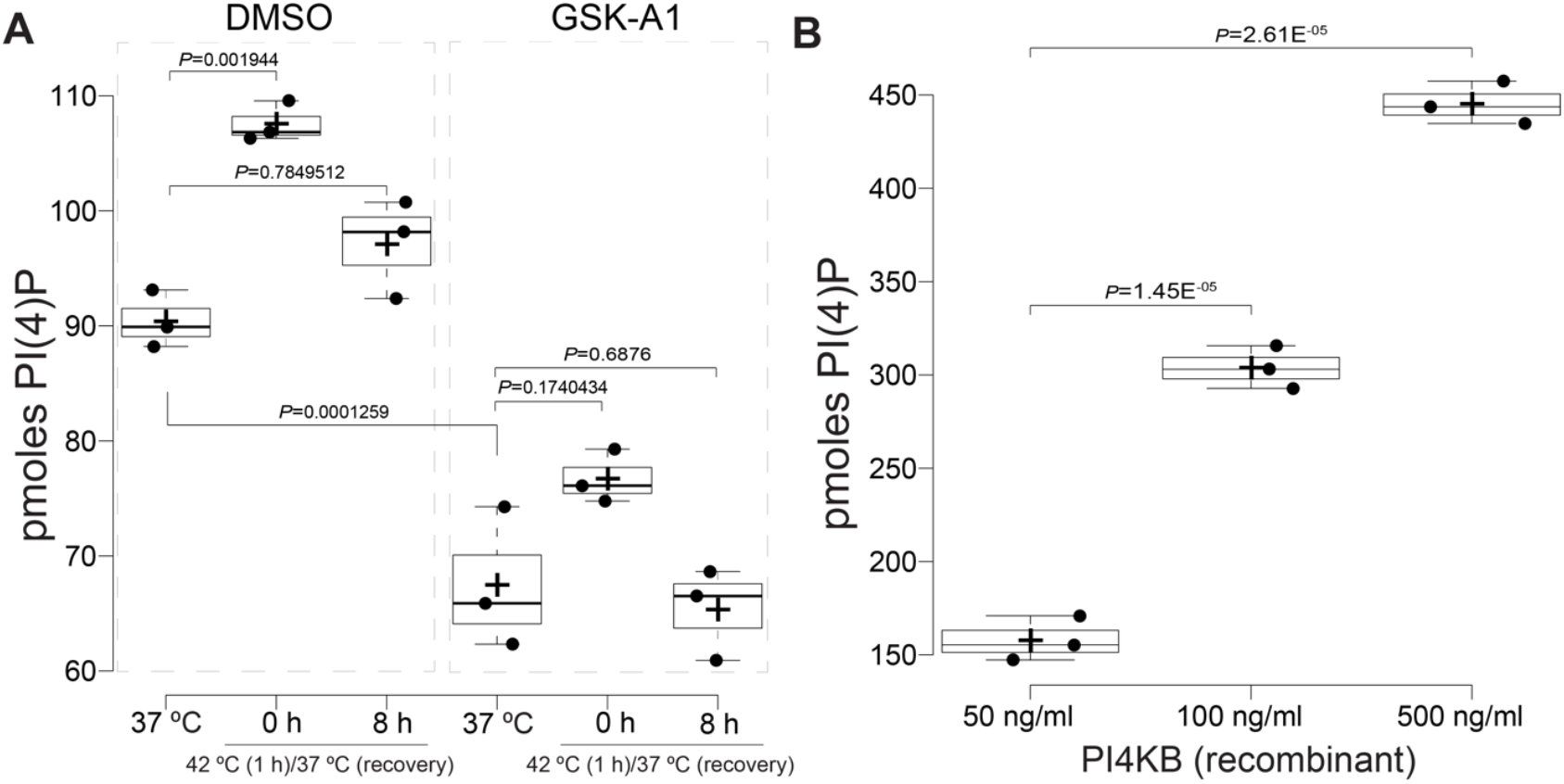
Heat shock increases PI4KIII alpha activity, which is suppressed by GSK-A1 treatment. (A) PI4KIII alpha activity was assessed using the PI4-Kinase Activity ELISA kit (Echelon Biosciences, Salt Lake, UT, USA). Enzyme activity significantly increased immediately after heat shock (0 h), followed by a decline during recovery at 8 h (left plots). Treatment with GSK-A1, a PI4K inhibitor, substantially reduced overall activity (right plots). Data represent three independent experiments performed on a separate batch of cells, with individual data points shown as closed circles. (B) Validation of the assay using recombinant PI4KB protein, demonstrating a concentration-dependent increase in PI4P production. For both A and B, center lines indicate medians, box limits represent the 25th and 75th percentiles, whiskers extend 1.5 times the interquartile range, and crosses denote sample means. Statistical significance was determined using one-way ANOVA followed by post-hoc Tukey HSD and Bonferroni tests.

These results suggest that the observed PI(4)P increases are primarily driven by enhanced PI4KIII alpha activity rather than changes in its expression, underscoring the role of post-translational regulation in the heat shock response.

### Dependency of HSPA1A Translocation on PI(4)P and PI4KIII Alpha

We have previously established the critical role of PI(4)P in facilitating the PM localization of HSPA1A (19). With the new findings that PI(4)P levels at the PM increase following heat shock (HS), and this increase depends on PI4KIII alpha activity, we sought to further confirm PI(4)P as a trigger for HSPA1A’s PM translocation.

Using RNA interference (RNAi) to deplete PI4KIII alpha, we observed a significant loss of the PI(4)P-specific biosensor SIDM from the PM at both 0 h and 8 h post-HS (Figure 8A and Supplement Figure 7), indicating that PI4KIII alpha is essential for maintaining PI(4)P levels. This depletion also resulted in a corresponding loss of HSPA1A localization at the PM, as visualized through confocal microscopy (Figure 8B and Supplement Figure 7).

**Figure 8.**
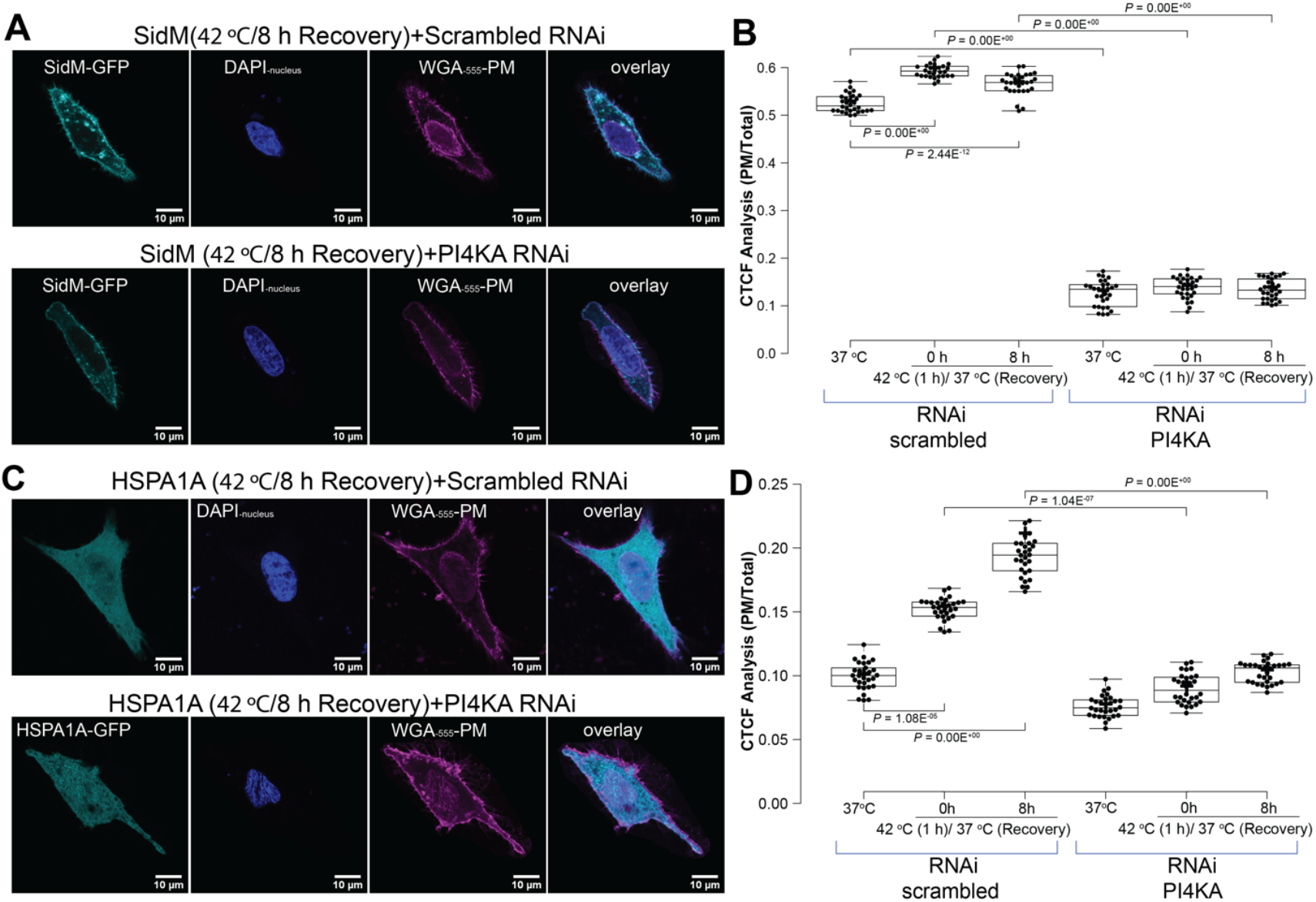
HSPA1A plasma membrane (PM) localization depends on PI4P and PI4KIII alpha activity. (A) Representative confocal images of HeLa cells showing the localization of the PI4P-specific biosensor SidM-P4M under control conditions (37 °C) immediately after heat shock (0 h recovery at 37 °C) and following 8 h of recovery. Cells were stained with WGA-FA555 (PM stain) and DAPI (nucleus stain) and transfected with either scrambled RNAi (top rows) or RNAi targeting PI4KIII alpha (bottom rows). PI4KIII alpha RNAi abolished SidM-P4M localization at the PM at 0 h and 8 h recovery. Scale bar = 10 μm. (B) Quantification of corrected total cell fluorescence (CTCF) as the ratio of SidM-P4M fluorescence at the PM to the rest of the cell. Data were collected under the conditions described in (A). Results are based on three independent experiments, with 30 cells per condition (closed circles). (C) Representative confocal images showing the effect of PI4KIII alpha RNAi on HSPA1A localization at the PM under the same conditions. Cells were transfected with scrambled RNAi (top rows) or RNAi targeting PI4KIII alpha (bottom rows). PI4KIII alpha depletion resulted in the loss of HSPA1A at the PM at both 0 h and 8 h recovery. Scale bar = 10 μm. (D) Quantification of the CTCF ratio of HSPA1A fluorescence at the PM to the rest of the cell. Data are based on three independent experiments representing total individual cell counts (N = 30). For both B and D, box limits indicate the 25th and 75th percentiles, whiskers extend 1.5 times the interquartile range, and crosses denote sample means. Statistical significance was determined using one-way ANOVA followed by Tukey HSD and Bonferroni tests. Additional images and RNAi validation are provided in Supplementary Figure 7.

To further validate these findings, we treated cells with GSK-A1, a pharmacological inhibitor of PI4K. We observed a significant reduction in HSPA1A PM localization at 0 h, coinciding with the significant increase in PI(4)P levels post-HS (Figure 9 and Supplement Figure 8).

**Figure 9.**
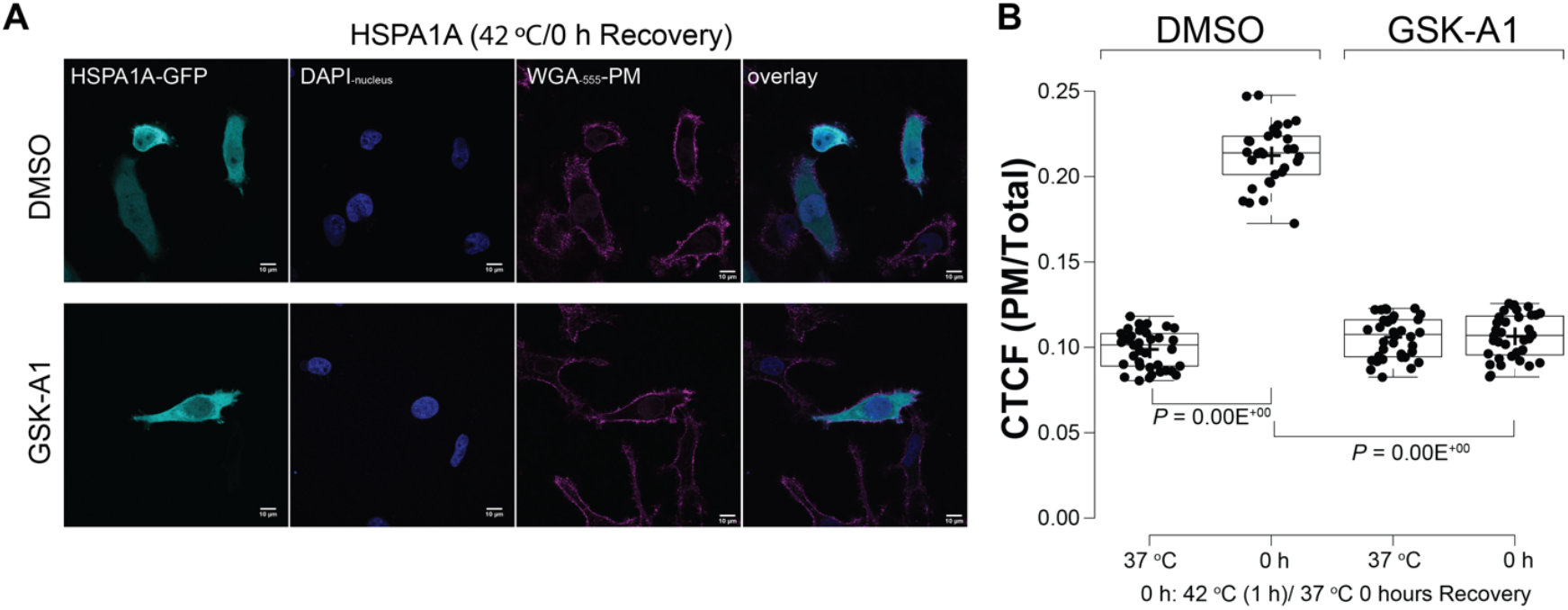
HSPA1A PM localization is inhibited in cells treated with GSK-A1, a PI4K inhibitor. (A) Representative confocal images of HeLa cells showing HSPA1A localization under control conditions (DMSO, top row) and following GSKA1 treatment (bottom row) at 0 h post-heat shock. Cells were stained with WGA-FA555 (PM stain) and DAPI (nucleus stain). Scale bar = 10 μm. (B) Quantification of HSPA1A localization at the plasma membrane (PM) in cells treated with DMSO or GSKA1, expressed as the corrected total cell fluorescence (CTCF) ratio of HSPA1A at the PM to the rest of the cell. Data represent three independent experiments, with total individual cell counts (N = 30; shown as closed circles). Center lines indicate medians; box limits represent the 25th and 75th percentiles; whiskers extend 1.5 times the interquartile range; crosses denote sample means. Statistical significance was determined using one-way ANOVA followed by Tukey HSD and Bonferroni tests. Additional images are provided in Supplementary Figure 8.

These results demonstrate the dependency of HSPA1A PM localization on PI(4)P levels and establish PI4KIII alpha activity as a key regulator of this process, highlighting the interconnected roles of lipid dynamics and kinase activity in stress-induced protein translocation.

## Discussion

This study identifies PI(4)P as a critical lipid mediator in the cellular stress response, demonstrating a significant increase in PI(4)P’s PM levels immediately following heat shock. This rapid PI(4)P enrichment is driven by enhanced PI4KIII alpha activity rather than transcriptional or translational changes. We further show that PI(4)P accumulation is essential for HSPA1A’s PM localization, establishing a mechanistic link between lipid signaling, kinase activity, and protein translocation (Figure 10). These findings underscore the pivotal role of lipid dynamics in the heat shock response. However, it is important to note that these findings are based on *in vitro* experiments using established cancer cell lines. While these models are invaluable for mechanistic studies, their lipid composition and stress responses may not fully recapitulate those of primary cells or tissues under physiological conditions.

**Figure 10.**
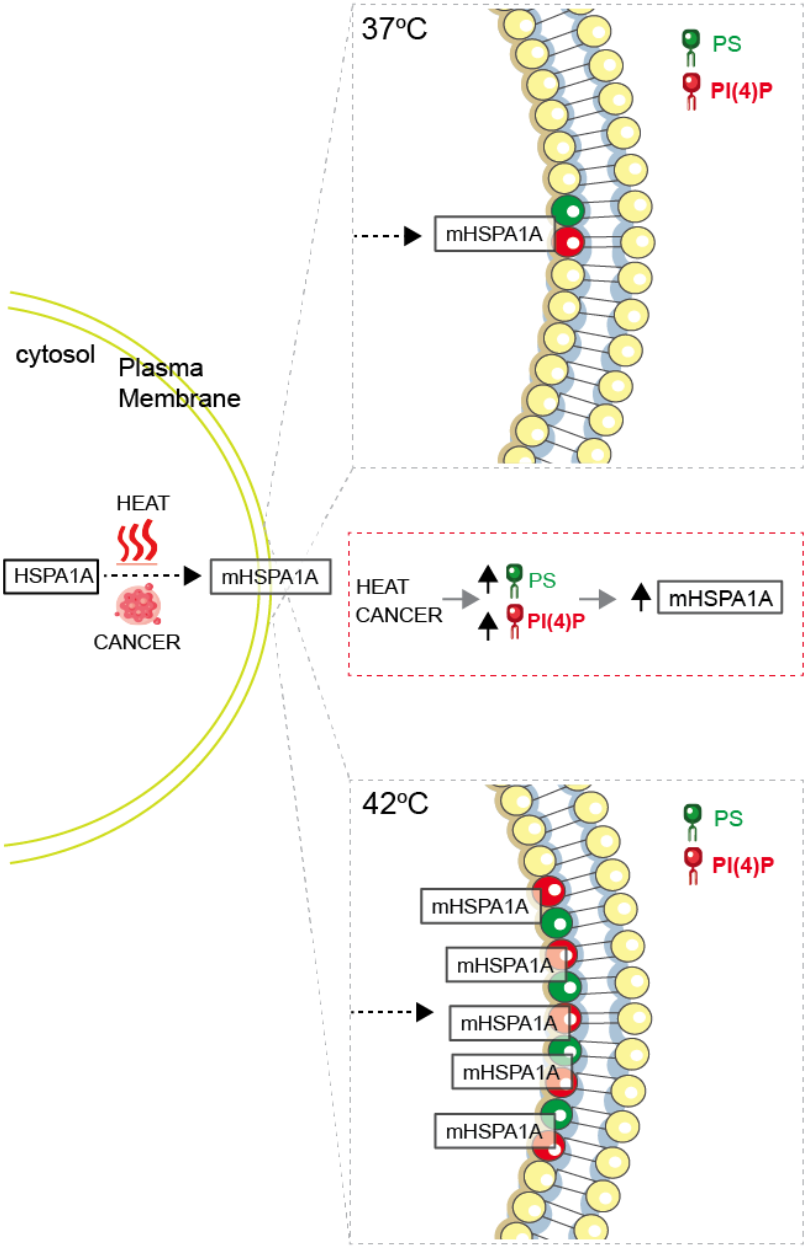
Model depicting the regulation of membrane HSPA1A (mHSPA1A) by specific lipids under stress conditions. At physiological temperature (37 °C), HSPA1A exhibits limited plasma membrane (PM) association, with phosphatidylserine (PS, green) and phosphatidylinositol 4-phosphate (PI(4)P, red) contributing to its baseline localization (top inset). During heat shock (42 °C) or in cancerous conditions, there is a significant increase in PS and PI(4)P levels at the PM (middle inset), correlating with enhanced HSPA1A translocation to the PM (bottom inset).

PI(4)P acts as a lipid mediator that enables HSPA1A’s recruitment to the PM, a process critical for maintaining membrane integrity and cellular homeostasis during stress (3,5,39,40). Unlike canonical lipid-binding proteins, HSPA1A’s PM localization is not merely passive but depends on PI(4)P enrichment triggered by heat shock. This discovery highlights a novel function for PI(4)P in stress-induced membrane remodeling. Furthermore, the dependence of HSPA1A translocation on PI(4)P provides a mechanistic explanation for the dynamic lipid changes observed in stressed and cancer cells, positioning PI(4)P as a key regulator of adaptive responses.

Our findings expand on previous studies highlighting the role of phosphoinositide dynamics in stress responses. The heat-induced increase in PI(4)P levels we observed parallels reports in yeast, where heat shock triggers rapid PI(4)P redistribution to the PM (33), and in plants, where heat stress enhances PI4K activity (32). These conserved responses across species emphasize PI(4) P’s universal role in maintaining membrane integrity and facilitating cellular adaptation to stress.

In addition to its role in HSPA1A translocation, PI(4)P likely influences a broader range of membrane dynamics, signaling, and trafficking processes under stress conditions. While Toker and Cantley (41) established the significance of PI3K-mediated signaling in stress adaptation, our results identify a distinct, rapid mechanism mediated by PI(4)P. This finding complements the phosphoinositide signaling network described by Balla (42), adding new insights into how lipid dynamics regulate stress-induced membrane processes.

Our data indicate that the rapid PI(4)P increase post-heat shock is driven by PI4KIII alpha activity rather than its expression, aligning with the heat shock response paradigm where gene expression is broadly suppressed except for heat shock proteins (37). The enhanced activity of PI4KIII alpha might involve heat-induced post-translational modifications (33), release from inhibitory complexes (31), increased substrate availability (PI increases during heat shock Figure 2 and (30,43), or structural changes in the PM that create PI-enriched microdomains.

These mechanisms warrant further investigation to uncover general principles of lipid-mediated stress adaptation.

The observed rapid activation of PI4KIII alpha may parallel mechanisms in yeast, where heat shock disrupts Osh3 lipid transfer protein function, leading to PI(4)P accumulation (33). Similarly, the interplay between PI(4)P and PI(4,5)P_2_ metabolism during stress, as documented in plants (32) and fungi (30), suggests that PI4KIII alpha activation is part of a broader phosphoinositide remodeling process to optimize membrane composition for stress tolerance.

Cancer cells leverage elevated stress tolerance and lipid remodeling to thrive in hostile microenvironments (24,44,45). Our findings highlight the dependency of HSPA1A translocation on PI(4)P (Figure 10) and PI4KIII alpha activity, underscoring their roles in stabilizing the PM, preventing cell lysis, and supporting pro-survival signaling pathways. These mechanisms likely contribute to tumor progression and therapeutic resistance. Targeting PI4KIII alpha or modulating PI(4)P levels with pharmacological inhibitors like GSK-A1 could disrupt these adaptive processes. Combining PI4KIII alpha inhibition with therapies targeting stress response pathways may yield synergistic effects, enhancing cancer cell sensitivity to treatment.

Beyond HSPA1A translocation, stress-induced PI(4)P accumulation may influence other cellular processes, such as stress fiber formation and altered motility, similar to PIP_2_ in other stress contexts (46). PI(4) P’s role in the Golgi stress response (47,48) suggests that targeting PI4KIII alpha could disrupt multiple stress adaptation pathways, broadening the therapeutic potential for combating cancer resilience.

Our previous work demonstrated that heat-induced increases in PS are essential for HSPA1A’s PM translocation (21). Expanding on this, we now show that PI(4)P also plays a critical role in facilitating HSPA1A’s membrane localization following heat shock. This suggests that multiple lipid signals cooperate to regulate HSPA1A translocation, highlighting a broader lipid-driven mechanism underlying the stress response. Given that PS and PI(4)P are both enriched at the PM and their levels increase in response to heat stress, it is possible that they function in a coordinated manner to stabilize membrane-associated HSPA1A. Future work examining whether PI(4)P facilitates HSPA1A’s initial recruitment to PS-enriched domains or whether these lipids act in parallel to reinforce membrane association would further elucidate the coordinated role of lipid signaling in stress adaptation and membrane remodeling.

While our findings shed light on the role of PI(4)P and PI4KIII alpha in stress responses, several questions remain unanswered. Specifically, the mechanisms that regulate PI4KIII alpha activity during heat shock are not fully understood. Identifying these regulatory pathways could enhance our understanding of lipid kinase regulation under stress. Additionally, while we’ve focused on HSPA1A, it’s unclear how PI(4)P interacts with other membrane-associated proteins, potentially uncovering broader roles for PI(4)P in cellular stress responses. The spatial and temporal dynamics of PI(4)P changes across cellular compartments also require further investigation. Finally, expanding this work to primary cells, including non-cancerous cells or *in vivo* models, will be essential to determine whether these mechanisms are universal or specific to cancer and heat shock and will be crucial for translating these findings into therapeutic strategies.

Our study highlights the critical role of PI(4)P and PI4KIII alpha activity in regulating HSPA1A translocation to the PM during the heat shock response (Figure 10). We provide new insights into the dynamic interplay between lipid signaling and stress adaptation by uncovering a lipid-mediated mechanism that operates independently of transcriptional or translational changes. This rapid lipid-driven response likely represents a fundamental strategy for maintaining membrane integrity and function under diverse conditions. These findings deepen our understanding of cellular resilience mechanisms and open new avenues for targeting stress response pathways in cancer therapy. Integrating these molecular insights with therapeutic strategies may pave the way for innovative treatments to disrupt lipid-regulated survival mechanisms in cancer and other stress-associated diseases.

## Materials and Methods

### Plasmids

To study HSPA1A localization, we utilized differentially tagged protein versions previously described in our earlier publications. Specifically, the mouse *hspa1a* cDNA sequence (accession number BC054782) was used to generate the recombinant constructs employed in this study. HSPA1A was tagged with green fluorescent protein (GFP). The gene was subcloned into the pEGFP-C2 vector (producing N-terminally tagged HSPA1A) for expression in mammalian cells. Subcloning was performed via directional cloning after PCR amplification and restriction enzyme digestion, following protocols detailed in the references (18) and (49). The PS biosensor Lact-C2-GFP plasmid was generously provided by Sergio Grinstein (Addgene plasmid #22852) (50). The PI(4)P-biosensor GFP-P4M-SidMx2 was a gift from Tamas Balla (Addgene plasmid # 51472) (51).

### Cell Culture

For all experiments described in this study, we utilized HeLa cells derived from Henrietta Lacks (ATCC® CCL-2™), obtained from ATCC in December 2016 and verified bi-annually. HeLa cells were grown in Minimum Essential Medium (MEM) supplemented with 10% fetal bovine serum (FBS), 2 mM L-glutamine, 0.1 mM non-essential amino acids (NEAA), 1 mM sodium pyruvate, and penicillin-streptomycin. All cultures were maintained in a humidified atmosphere containing 5% CO_2_ at 37 °C. Cells were grown into 24-well plates (2.0 × 10^4^ cells per well), 60mm^2^ plates (3.0 x 10^6^ cells per plate), or 75 cm^2^ cell culture flasks (6.0 × 10^6^ cells per flask).

### Transient Cell Transfections

#### Plasmid Transfections

Transient transfections were performed to express the proteins used in this study. Cells were seeded one day before transfection into 24-well plates (2.0 × 10^4^ cells per well, with poly-D-lysine-treated coverslips). After 18 hours, cells were transfected with the appropriate construct using the PolyJet In Vitro DNA Transfection Reagent (SignaGen Laboratories, Frederick, MD, USA), following the manufacturer’s instructions. Transfection proceeded for 18 hours, after which the media was replaced with fresh, complete media to support subsequent experiments.

#### RNAi Transfections

To investigate the role of PI(4)P synthesis enzyme in HSPA1A’s PM localization, RNA interference (RNAi) was used to inhibit PI4KA. The impact of these genetic interventions on HSPA1A localization and protein levels was assessed using confocal microscopy. Non-targeting control siRNA (Stealth^TM^ RNAi Negative control duplexes, Cat. No. 12935-300; Invitrogen, Carlsbad, CA, USA) was used as a negative control to account for off-target effects of RNAi.

For all RNAi transfections, cells were co-transfected with 0.75 µg of DNA encoding HSPA1A-GFP or SidM-GFP. Experimental cells were transfected with 1.50 pmol of PI4KA RNAi (PI4KA Stealth siRNA, Cat. No. 1299001, assay ID: HSS108026; Invitrogen, Carlsbad, CA, USA), while control cells received 0.25 nM of non-targeting control siRNA. Transfections were carried out using the Lipofectamine™ 3000 Transfection Reagent, prepared by incubating the RNAi/DNA mixture with the reagent for 15 minutes at room temperature. This mixture was added to cells seeded on poly-D-lysine-coated coverslips in 24-well plates. Transfection proceeded for 24 hours, followed by the replacement of transfection media with fresh, complete media.

### Cell Treatments

*Heat Shock Treatment:* To assess the effect of heat shock on HSPA1A relocalization and cellular lipids, cells were either maintained at 37 °C or subjected to heat stress by incubating in a humidified CO_2_ incubator equilibrated at 42 °C for 60 minutes. Following heat shock, cells were used directly (0 hours of recovery; 0 h) or recovered at 37 °C for 8 hours (8 h).

#### PS Inhibition

To investigate the impact of plasma membrane PS reduction on SidM and HSPA1A PM localization, cells were treated with fendiline (Cayman Chemical, Ann Arbor, MI, USA), an FDA-approved drug known to decrease PS content in MDCK (52) and HEK293 cells (53). Cells were exposed to either DMSO (control) or 10 µM fendiline diluted in serum-free MEM for 48 hours at 37 °C in a humidified CO_2_ incubator. Fendiline was added directly to the existing culture media without prior aspiration. Heat shock and recovery protocols were synchronized to conclude at the end of the 48-hour treatment.

#### PI(4)P Inhibition

To investigate the impact of PI(4)P reduction on SidM and HSPA1A PM localization cells were treated with GSK-A1 (AdipoGen Life Sciences, San Diego, CA, USA), a specific inhibitor of PI4KA (54). Cells were exposed to either DMSO (control) or 100 nM GSK-A1 diluted in serum-free MEM for 30 minutes at 37 °C in a humidified CO_2_incubator. The compound was added directly to the existing culture media without prior aspiration. Heat shock and recovery protocols were synchronized to conclude at the end of the 30-minute treatment.

#### Cell Viability Assay

Cell viability following heat shock and other treatments was assessed using the trypan blue exclusion assay. Viable and non-viable cells were quantified with the Cellometer® Auto X4 Cell Counter (Nexcelom Bioscience, Lawrence, MA, USA).

### Confocal Microscopy and Image Analysis

#### Cell Fixation and Staining

After transfection and subsequent treatments, cells were fixed in 4% paraformaldehyde (PFA) prepared in a complete growth medium for 12 minutes at room temperature (RT). Following fixation, PFA was aspirated, and the cells were washed three times with 1X phosphate-buffered saline (PBS). To stain the PM, cells were incubated with 1 μg/ml wheat germ agglutinin (WGA) Alexa Fluor® 555 conjugate (Invitrogen, Carlsbad, CA, USA for 10 minutes at RT. After staining, the cells were washed three additional times with 1X PBS. Coverslips were mounted onto slides using approximately 3 μL of DAPI Fluoromount-G® (SouthernBiotech, Birmingham, AL, USA) and allowed to dry for 24 hours at RT in the dark. Prepared slides were stored at 4 °C until visualization by confocal microscopy (19).

#### Microscope Setup

Cell imaging was performed using an Olympus FLUOVIEW FV3000 inverted scanning confocal microscope with a 60x 1.5 NA oil immersion objective. Multichannel image acquisition was conducted with the following settings: Channel 1 (Green): Excitation at 488 nm; emission at 510 nm; Channel 2 (Blue): Excitation at 405 nm; emission at 461 nm; Channel 3 (Red): Excitation at 561 nm; emission at 583 nm.

#### Image Analysis

Quantitative analysis was conducted on images from several cells (details provided in figure legends) collected across three independent experiments. Images were processed manually using ImageJ software (55), employing the corrected total cell fluorescence (CTCF) method (56). The formula used for CTCF was: CTCF = Integrated Density – (Area of Region of Interest * Fluorescence of background reading).

CTCF values were determined for both the PM and the cytosol. A ratio of PM fluorescence to total cell fluorescence (PM + cytosol) was calculated for each cell, following established protocols (49,57-59). As cell slices are not entirely flat, and the CTCF method utilizes the outermost pixels of each cell, a PM localization value of 0% cannot be achieved (57,58).

## Western Blot Analysis

For western blot analysis, 15 μg of total cell lysate was used for total protein detection. The following antibodies were used:

- Anti-HSP70 Monoclonal Antibody (Mouse IgG; clone C92F3A-5; dilution 1:1000), Enzo Life Sciences (Farmingdale, NY, USA).
- Anti-Na+/K+ ATPase α (ATP1A1) Antibody RabMAb® (EP1845Y; 2047-1; dilution 1:1000), Thermo Fisher Scientific (St. Louis, MO, USA).
- Anti-PI4KA Polyclonal Antibody (Rabbit IgG; 12411-1-AP; dilution 1:500), Thermo Fisher Scientific (St. Louis, MO, USA).
- Anti-GAPDH Monoclonal Antibody (Mouse IgG2b; 60004-1-Ig; dilution 1:1000), Proteintech Group, Inc (Rosemont, IL, USA)

Blots were incubated with primary antibodies overnight (∼16 hours) at 4 °C with constant rotation. Total protein staining was performed using Pierce™ Reversible Protein Stain (Thermo Scientific Pierce, Rockford, IL, USA).

*Signal Detection:* Western blot signals were visualized using the Omega Lum™ C Imaging System (Aplegen, San Francisco, CA).

### Lipid Quantification and Analysis

#### Mass Spectrometry and Lipidome Analysis

##### Sample Preparation

Six biological replicates (different cell batches) of four million HeLa cells were prepared to identify total lipid profiles before and after heat shock. Samples were collected under three conditions: control (37 °C), immediately after heat shock (0 h), and after an 8-hour recovery period following heat shock (8 h).

##### Lipid Extraction and Analysis

Samples were analyzed by the UC Davis West Coast Metabolomics Center using biphasic lipid extraction, followed by data acquisition on a C18-based hybrid bridged column with a ternary water/acetonitrile/isopropanol gradient and a ThermoFisher Scientific Q-Exactive HF mass spectrometer with electrospray ionization. MS/MS data were acquired in data-dependent mode, and accurate masses were normalized by constant reference ion infusion. MS-DIAL vs. 4.90 was used for data processing and lipid annotations (60,61). Lipids were annotated in identity search mode with precursor mass errors < 10 mDa, retention time matching to the corresponding lipid classes, and MS/MS matching. Data were sum-normalized: each lipid was represented as a fraction of total lipids in the sample and scaled by the median intensity of all lipids within the treatment group (62-64). Data are shown in Supplementary Table 1. Heatmap analysis (MetaboAnalyst (34)) was performed to simultaneously visualize lipidomic changes in response to experimental conditions and cluster similar samples based on their lipid profiles. Total lipid abundance was visualized using boxplot graphs (65).

##### PI(4)P Mass ELISA Assay

To quantify PI(4)P levels after heat shock, we used the PI(4)P Mass ELISA kit (K-4000E; Echelon Biosciences, Salt Lake, UT, USA), following the manufacturer’s protocol. Briefly, standards and PI(4)P samples were prepared by resuspension in PBS-T buffer, followed by sonication and incubation with the PI(4)P detector and secondary antibodies. After incubation with TMB substrate, absorbance was measured at 450 nm to generate a standard curve and calculate PI(4)P levels.

Lipids were extracted using a modified MeOH:CHCl_3_ protocol. Briefly, 2 or 4 million cells were treated with 0.5 M TCA to precipitate cellular contents. Neutral lipids were removed with MeOH:CHCl_3_ (2:1), and acidic lipids, including PI(4)P, were extracted with MeOH:CHCl_3_:HCl (80:40:1). The organic phase was isolated, dried under vacuum, and stored at -20 °C until use.

### Enzyme Activity Assay for PI4KIII Alpha

To assess PI4KIII alpha activity, we utilized the PI4-Kinase Activity ELISA kit (K-4000K; Echelon Biosciences, Salt Lake, UT, USA), following the manufacturer’s protocol. The assay is a competitive ELISA that detects and quantifies PI(4)P produced from a PI4-Kinase reaction.

Briefly, PI substrate and ATP were prepared, and enzyme reactions were initiated in the presence of specific buffers. After incubation, reactions were stopped with EDTA, and the resulting PI(4)P was quantified using a PI(4)P Detector and colorimetric detection at 450 nm. PI4K activity was interpolated from a standard curve generated with serial PI(4)P standard dilutions.

For immunoprecipitation of PI4KIII alpha, cell lysates were prepared from 3×10^6^ cells using ice-cold lysis buffer (25 mM Tris pH 8.0, 150 mM NaCl, 1% NP-40, 1 mM EDTA, 5% Glycerol) with protease inhibitors. PI4KA was immunoprecipitated with the PI4KA Polyclonal Antibody (12411-1-AP; dilution 1:500; Thermo Fisher Scientific, St. Louis, MO, USA) overnight at 4 °C under constant rotation. The mixture was then incubated for 3 hours with A/G agarose beads (Pierce™ Protein A/G Agarose; catalog number: 20422; Thermo Scientific Pierce, Rockford, IL, USA) at 4 °C under constant rotation. After extensive washing, the bead-enzyme complexes were directly used in the PI4-Kinase Activity Assay (10 μl per well).

### RNA Sequencing and Transcript Analysis

Cells from the control, 0R, and 8R conditions were sent to Novogene (Sacramento, CA) for RNA isolation, library preparation, and sequencing. Briefly, total RNA from control, 0R, and 8R conditions was isolated and quality-checked before cDNA library preparation and paired-end sequencing on an Illumina NovaSeq platform, yielding ≥30 million reads per sample. Reads underwent adapter trimming, PhiX filtering, and quality control using BBDuk. High-quality reads were aligned to the human reference genome (GRCh38.p13) with STAR, generating splice-aware BAM files. Mapped reads were sorted and indexed with Samtools for feature counting using HTSeq, producing raw gene counts. Quality control metrics were summarized with MultiQC, and differentially expressed genes were identified using DESeq2. The complete methods, data, and analyses are described in (38) and the raw data are hosted at NCBI (GEO: GSE285497).

*qPCR Validation of PI4 Kinase Transcripts:* Following the manufacturer’s protocol, RNA was isolated using three million HeLa cells per condition (control cells, 0-hour recovery, 8-hour recovery; different batches from the ones used for RNA-seq) using the Direct-Zol RNA mini-prep Kit (ZymoResearch, Irvine, CA, USA). Following the manufacturer’s protocol, cDNA was synthesized from 1ug of total RNA using the Superscript IV First-Strand synthesis system (ThermoFisher Scientific) and Oligo(dT)_20_ primers. cDNA samples were diluted to a concentration of 50 ng/uL.

Three biological replicates were used for each gene and condition [Gene names and primers (generated using NCBI’s primer-blast utility) are shown in Supplementary Table 3]. For qPCR, reactions were prepared in a 20 μL total volume using Power SYBR™ Green PCR Master Mix (ThermoFisher Scientific, Waltham, MA, USA). Each reaction contained 1 μL cDNA (10 ng), 0.6 μL of each primer (300nM), and 10 μL SYBR Green mix. Amplification was performed on a CFX96 Touch Real-Time Detection System (Bio-Rad, Hercules, CA, United States) with the following thermal cycling conditions: initial denaturation at 95 °C for 10 minutes, followed by 45 cycles of denaturation at 94 °C for 10 seconds, annealing at 50 °C for 10 seconds, and extension at 68 °C for 10 seconds. The relative normalized expression (66) of the raw transcript levels was calculated using the Livak method for each gene (67) using the software provided with the instrument (68). The reference genes used in this method were ACTB and GAPDH.

For endpoint RT-PCR, 100 ng of cDNA was used per reaction. The PCR reaction (20 μL total volume) contained 0.5 μL of each primer (10 μM), dNTP (10 mM), and 0.5 units of Taq polymerase. The amplification protocol consisted of 40 cycles under the following conditions: an initial denaturation at 95 °C for 5 minutes, followed by denaturation at 94 °C for 30 seconds, annealing at 50 °C for 20 seconds, and extension at 72 °C for 20 seconds, with a final extension step at 72 °C for 5 minutes. PCR products were analyzed on a 2% agarose gel, stained with ethidium bromide, and visualized under UV light.

### Statistical Methods

Statistical significance was assessed using one-way ANOVA (Analysis of Variance) followed by post-hoc Tukey HSD (Honestly Significant Difference) and Bonferroni tests. A *P* value < 0.05 was considered statistically significant. Boxplots were generated using the BoxPlotR application (http://shiny.chemgrid.org/boxplotr/) (65).

## Abbreviations

ANOVA: Analysis of Variance
cDNA: Complementary DNA
CTCF: Corrected Total Cell Fluorescence
DMSO: Dimethyl Sulfoxide
DMEM: Dulbecco’s Modified Eagle Medium
ELISA: Enzyme-Linked Immunosorbent Assay
FBS: Fetal Bovine Serum
GFP: Green Fluorescent Protein
HSD: Honestly Significant Difference
HSPA1A: Heat Shock Protein A1A
LDS: Lithium Dodecyl Sulfate
MEM: Minimum Essential Medium
MS/MS: Tandem Mass Spectrometry
NEAA: Non-Essential Amino Acids
PBS: Phosphate-Buffered Saline
PBS-T: Phosphate-Buffered Saline-TritonX100
PFA: Paraformaldehyde
PI4KA: Phosphatidylinositol 4 Kinase III alpha
PI(4)P: Phosphatidylinositol 4-Phosphate
PM: Plasma Membrane
PS: Phosphatidylserine
qPCR: Quantitative Polymerase Chain Reaction
QTOF: Quadrupole Time-of-Flight
RNAi: RNA Interference
RT: Room Temperature
SDS-PAGE: Sodium Dodecyl Sulfate-Polyacrylamide Gel Electrophoresis
siRNA: Small Interfering RNA
TMB: 3, 3’, 5, 5’ – Tetramethylbenzidine
WGA: Wheat Germ Agglutinin.

