## supplemental figures for "Heat Shock-Induced PI(4)P Increase Drives HSPA1A Translocation to the Plasma Membrane in Cancer and Stressed Cells through PI4KIII Alpha Activation"

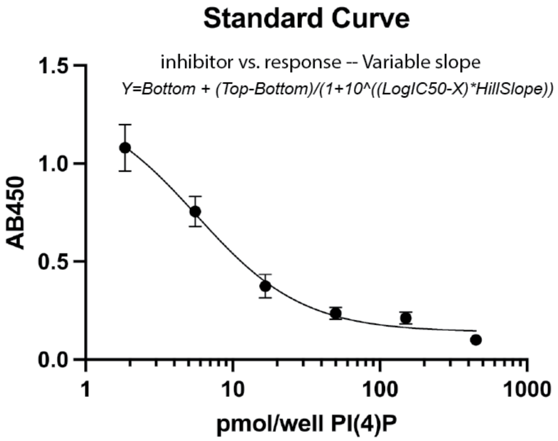


| Best-fit values ($R^2$=0.9696) | |
|---|---|
| Bottom | 0.1410 |
| Top | 1.329 |
| IC50 | 5.683 |
| HillSlope | -1.202 |

**Supplemental Figure 1. PI(4)P Standard Curve and Best-Fit Values.** Left panel: PI(4)P standard curve generated using non-linear regression analysis with GraphPad Prism version 10. The analysis used an agonist vs. response — Variable slope (four parameters) method. Right panel: Best-fit values derived from the standard curve analysis.

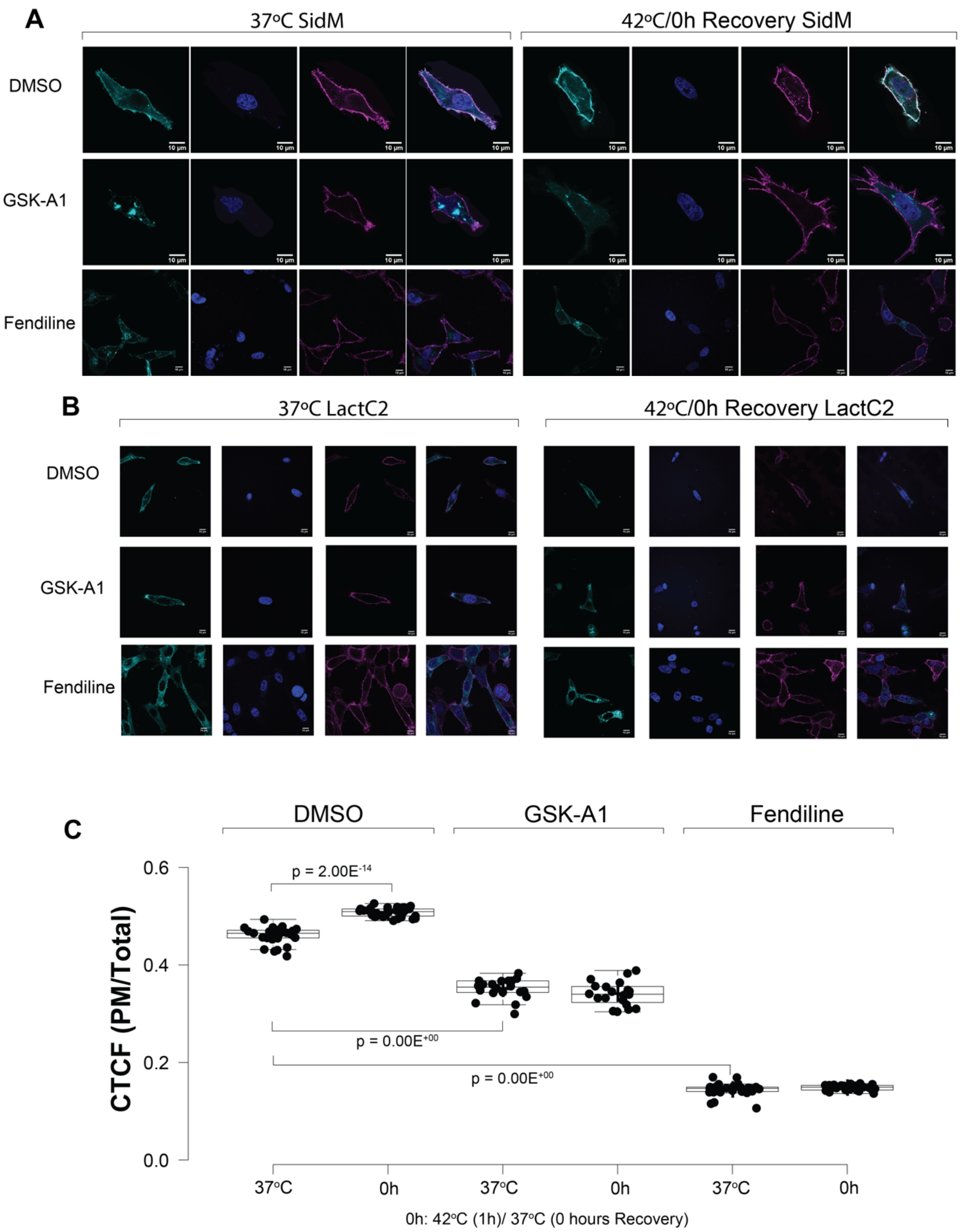


**Supplemental Figure 2.** Heat shock increases PI(4)P and phosphatidylserine (PS) levels at the plasma membrane (PM), as shown by SidM-P4M and Lact-C2 localization. (A) Additional confocal images of HeLa cells expressing SidM-P4M-GFP, a PI4P-specific biosensor, shown in Figure 3. Cells were stained with WGA-FA555 to label the PM and DAPI to label nuclei. Images depict SidM-P4M localization under control conditions (37°C; left panels) and immediately following heat shock (1 h at 42°C; no recovery; right panels) using DMSO (top rows), GSK-A1 (middle rows), and Fendiline (bottom rows). Scale bar = 10 µm. (B) Representative confocal images of HeLa cells expressing GFP-Lact-C2, a PS biosensor. Cells were stained with WGA-FA555 to label the PM and DAPI to label nuclei. Images depict SidM-P4M localization under control conditions (37°C; left panels) and immediately following heat shock (1 h at 42°C; no recovery; right panels) using DMSO (top rows), GSK-A1 (middle rows), and Fendiline (bottom rows). Scale bar = 10 μm. (C) The graph shows the quantification of corrected total cell fluorescence (CTCF) as the ratio of Lact-C2-GFP fluorescence at the PM to the rest of the cell. Results are based on three independent experiments, with 30 cells per condition (closed circles). Box limits indicate the 25th and 75th percentiles, whiskers extend 1.5 times the interquartile range, and crosses denote sample means. Statistical significance was determined using one-way ANOVA followed by Tukey HSD and Bonferroni tests.

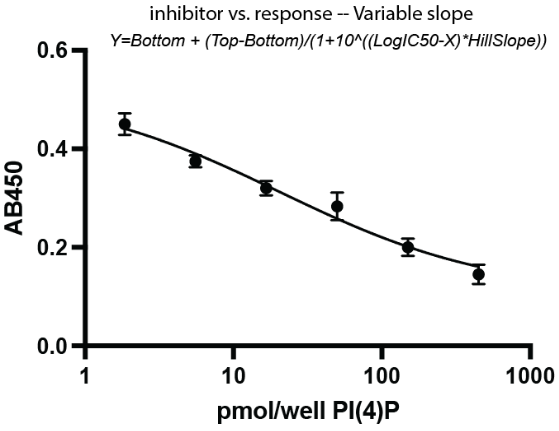


| Best-fit values ($R^2$=0.9661) | |
|---|---|
| Bottom | 0.1041 |
| Top | 0.5213 |
| IC50 | 20.48 |
| HillSlope | -0.5980 |

**Supplemental Figure 3. PI(4)P Standard Curve and Best-Fit Values.** Left panel: PI(4)P standard curve generated using non-linear regression analysis with GraphPad Prism version 10. The analysis was performed using an agonist vs. response — Variable slope (four parameters) method. Right panel: Best-fit values derived from the standard curve analysis.

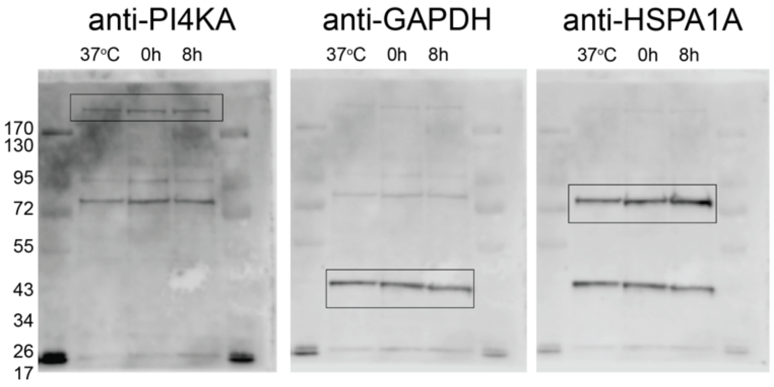


**Supplemental Figure 4.** Complete Western blots showing PI4KA protein amounts during recovery from mild heat shock. The uncropped Western blots correspond to the western blot experiments shown in Figure 6B. The blots were probed with PI4KA, GAPDH, and HSPA1A antibodies from left to right. Molecular size markers (Fisher BioReagents™ EZ-Run™ Prestained Rec Protein Ladder) are indicated for each blot.

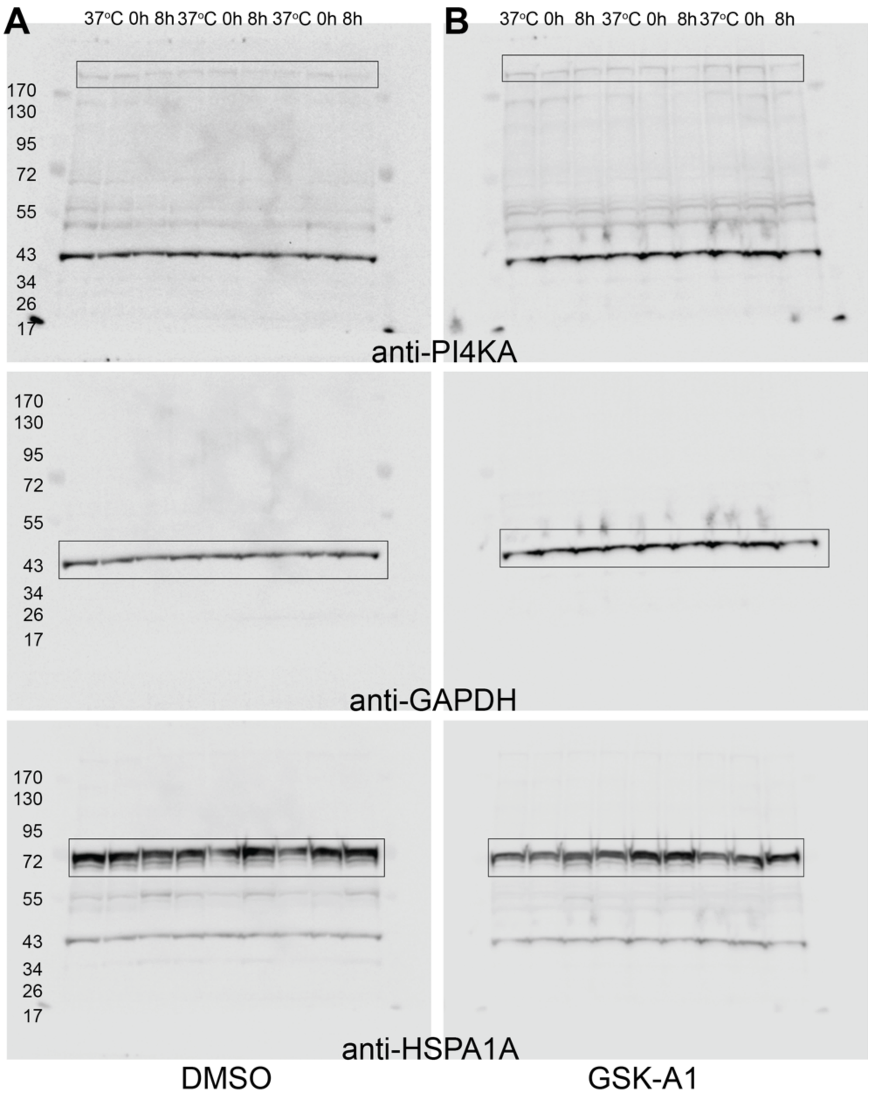


**Supplemental Figure 5. Input Samples for PI4KA Co-Immunoprecipitation used in the PI4-Kinase Activity Assay.** Western blot analysis of total protein lysates used as input for PI4KA co-immunoprecipitation (co-IP) experiments used in the ELISA assays shown in Figure 7A. Cells were treated with either DMSO (A) or the PI4KIIIα inhibitor GSK-A1 (B) and collected under three conditions: control (37°C), immediately after heat shock (0 h), and following an 8-hour recovery (8 h). These total lysates were used for immunoprecipitation but do not represent the co-immunoprecipitated proteins, since the immunoprecipitated material was used for the PI4-Kinase Activity ELISA. The blots were probed with anti-PI4KA to detect PI4KIIIα in total lysates before IP, anti-GAPDH as a loading control, and anti-HSPA1A as a positive control to confirm heat shock-induced upregulation. Molecular size markers (Fisher BioReagents™ EZ-Run™ Prestained Rec Protein Ladder) are indicated on the left of each blot.

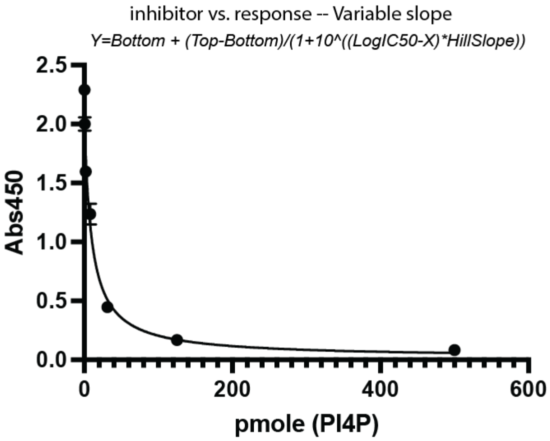


| Best-fit values ($R^2$=0.9807) | |
|---|---|
| Bottom | 0.01593 |
| Top | 2.053 |
| IC50 | 9863 |
| HillSlope | -0.9783 |

**Supplemental Figure 6. PI(4)P Standard Curve and Best-Fit Values.** Left panel: PI(4)P standard curve generated using non-linear regression analysis with GraphPad Prism version 10. The analysis was performed using an agonist vs. response — Variable slope (four parameters) method. Right panel: Best-fit values derived from the standard curve analysis.

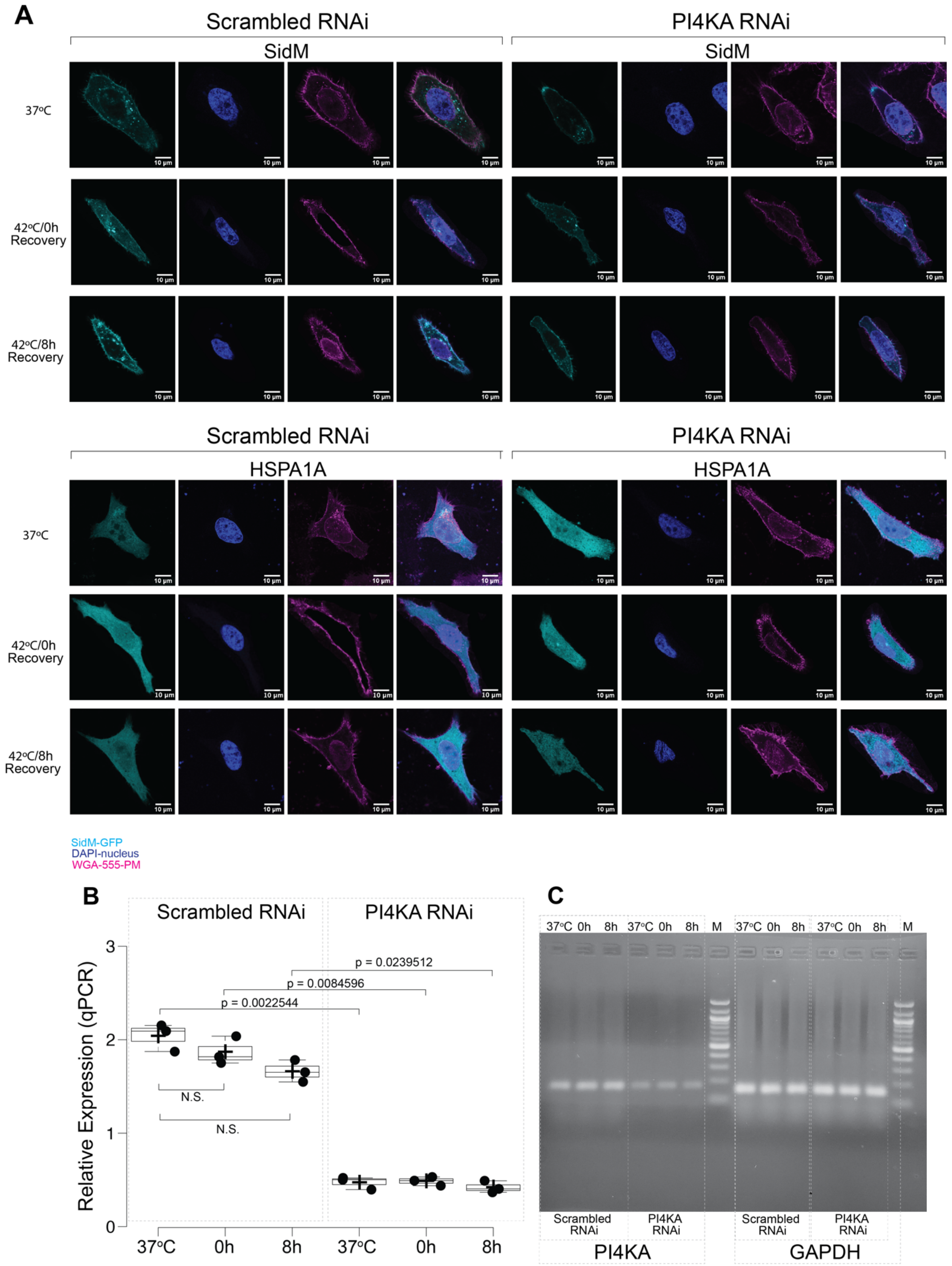
A
Scrambled RNAi
PI4KA RNAi
SidM
37ºC
42ºC/0h Recovery
42ºC/8h Recovery
HSPA1A
10 µm
SidM-GFP
DAPI-nucleus
WGA-555-PM
B
Scrambled RNAi
PI4KA RNAi
Relative Expression (qPCR)
p = 0.0239512
p = 0.0084596
p = 0.0022544
N.S.
N.S.
37ºC
0h
8h
C
37ºC 0h 8h 37ºC 0h 8h M 37ºC 0h 8h 37ºC 0h 8h M
Scrambled RNAi
PI4KA RNAi
PI4KA
GAPDH

**Supplemental Figure 7. HSPA1A plasma membrane (PM) localization depends on PI4P and PI4KIII alpha activity.** (A) Additional confocal images of HeLa cells showing the localization of SidM-P4M (top panels) and HSPA1A (bottom panels) under control conditions (37°C; top rows) immediately after heat shock (0 h recovery at 37°C; middle rows) and following 8 h of recovery (bottom rows). Cells were stained with WGA-FA555 (PM stain) and DAPI (nucleus stain) and transfected with either scrambled RNAi (left panels) or RNAi targeting PI4KIII alpha (right panels). Scale bar = 10 μm. (B) qPCR analysis of PI4KA transcripts using scrambled RNAi or RNAi targeting PI4III alpha. (C) RT-PCR analysis of PI4KA and GAPDH transcripts using scrambled RNAi or RNAi targeting PI4III alpha. M=100bp Ladder.

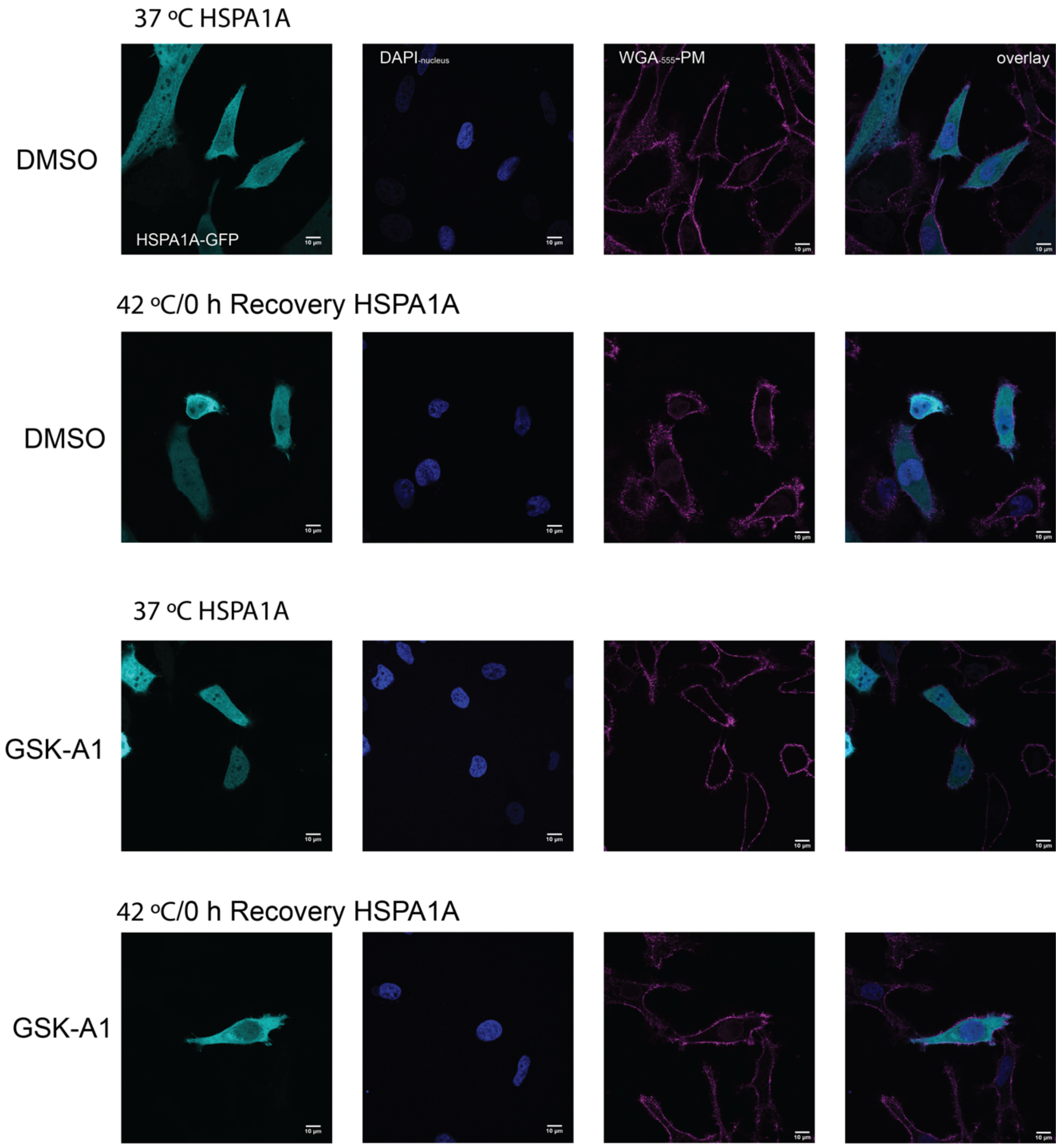


**Supplemental Figure 8. HSPA1A PM localization is inhibited in cells treated with GSK-A1, a PI4K inhibitor.** Additional confocal images of HeLa cells expressing HSPA1A, shown in Figure 9, under control conditions (DMSO, top rows) and following GSK-A1 treatment (bottom rows) at 37°C and 0 h post-heat shock. Cells were stained with WGA-FA555 (PM stain) and DAPI (nucleus stain). Scale bar = 10 μm.
