## supplemental Table 3 for "Heat Shock-Induced PI(4)P Increase Drives HSPA1A Translocation to the Plasma Membrane in Cancer and Stressed Cells through PI4KIII Alpha Activation"

**Supplementary Table 3.** Primers used in qPCR and RT-PCR experiments.

| Accession number | HGNC Symbol | Primer Sequence (5’-> 3’) | Primer Sequence (3’-5’) | Product Size (bp) |
| --- | --- | --- | --- | --- |
| NM_058004.4 | PI4KA | CAATCAAGCTCTTGAAGCACAG | CCTCGAAGGTCCCCTCCTC | 169 |
| NM_005345.6 | HSPA1A | AGCTGGAGCAGGTGTGTAAC | CAGCAATCTTGGAAAGGCCC | 154 |
| NM_001101.5 | ACTB | CTTCGCGGGCGACGAT | CCACATAGGAATCCTTCTGACC | 104 |
| NM_001256799.3 | GAPDH | CGGGAAGGAAATGAATGGGC | GGAAAAGCATCACCCGGAGG | 148 |
